# Endogenous CD28 drives CAR T cell responses in multiple myeloma

**DOI:** 10.1101/2024.03.21.586084

**Authors:** Mackenzie M. Lieberman, Jason H. Tong, Nkechi U. Odukwe, Colin A. Chavel, Terence J. Purdon, Rebecca Burchett, Bryan M. Gillard, Craig M. Brackett, A. J. Robert McGray, Jonathan L. Bramson, Renier J. Brentjens, Kelvin P. Lee, Scott H. Olejniczak

## Abstract

Recent FDA approvals of chimeric antigen receptor (CAR) T cell therapy for multiple myeloma (MM) have reshaped the therapeutic landscape for this incurable cancer. In pivotal clinical trials B cell maturation antigen (BCMA) targeted, 4-1BB co-stimulated (BBζ) CAR T cells dramatically outperformed standard-of-care chemotherapy, yet most patients experienced MM relapse within two years of therapy, underscoring the need to improve CAR T cell efficacy in MM. We set out to determine if inhibition of MM bone marrow microenvironment (BME) survival signaling could increase sensitivity to CAR T cells. In contrast to expectations, blocking the CD28 MM survival signal with abatacept (CTLA4-Ig) accelerated disease relapse following CAR T therapy in preclinical models, potentially due to blocking CD28 signaling in CAR T cells. Knockout studies confirmed that endogenous CD28 expressed on BBζ CAR T cells drove *in vivo* anti-MM activity. Mechanistically, CD28 reprogrammed mitochondrial metabolism to maintain redox balance and CAR T cell proliferation in the MM BME. Transient CD28 inhibition with abatacept restrained rapid BBζ CAR T cell expansion and limited inflammatory cytokines in the MM BME without significantly affecting long-term survival of treated mice. Overall, data directly demonstrate a need for CD28 signaling for sustained *in vivo* function of CAR T cells and indicate that transient CD28 blockade could reduce cytokine release and associated toxicities.

## Introduction

Chimeric Antigen Receptor (CAR) T cells are a form of immunotherapy that has seen extraordinarily success in treating hematologic malignancies^1–12^. Expression of a CAR in autologous T cells isolated from cancer patients redirects T cell specificity toward an antigen of interest and delivers activation signals upon antigen ligation^13–15^. Activation signals in clinically relevant second-generation CAR T cells are mediated by CD3ζ as well as a co-stimulatory domain, most commonly CD28 or 4-1BB, although others including ICOS and OX40 remain under investigation ^16–21^. Second generation CAR T cells targeting the B cell antigen, CD19, are FDA approved for B cell leukemias and lymphomas and have resulted in potent, decade long remissions for some of the initial patients who received this therapy^7,10,22–24^. Within the last few years, FDA approval for CAR T cells has expanded to include those directed against the tumor necrosis factor receptor (TNF-R) superfamily molecule, B cell maturation antigen (BCMA) for the treatment of multiple myeloma (MM)^25,26^. Despite robust initial response rates greatly outperforming standard of care in heavily pre-treated MM patient populations, recent clinical studies have shown that approximately 40% of MM patients will experience disease progression within the first 24 months of BCMA-targeted CAR T cell infusion ^27–29^. Therefore, there is an urgent need to understand and overcome resistance mechanisms that hinder CAR T cell success in MM patients.

MM remains an incurable malignancy of plasma cells, a terminally differentiated B cell subset that typically reside in the bone marrow (BM) and contribute to protective humoral immunity through the production of immunoglobulin. The long-term survival of plasma cells is critically dependent upon interactions occurring within the BM niche, with many soluble and contact-dependent stromal interactions also contributing to survival and disease progression of MM^30^. MM relies on CXCL12 chemokine gradients to home into the BM niche, as well as adhesion molecules including LFA-1 to mediate attachment and retention within the microvasculature^31–34^. Several soluble mediators secreted by BM resident dendritic cells (DCs), macrophages, osteoblasts, and stromal cells, including IL-6 and BCMA ligands, APRIL, and BAFF, sustain MM survival and proliferation^35–40^. Additionally contact-dependent interactions regulate anti-apoptotic molecule expression and chemotherapeutic resistance in MM. A key mediator of contact-dependent survival in MM is the canonical T cell co-stimulatory receptor, CD28, whose expression on MM cells is highly correlated with myeloma progression^41^. Importantly, CD28 ligands CD80 and CD86 are expressed on DCs, stromal cells, and even MM cells within the BM microenvironment^30,42–44^. Ligation of CD28 on MM cells by CD80/CD86 transduces a PI3K/Akt pathway dependent, pro-survival signal protecting them from chemotherapy and growth factor withdrawal-induced death^42,45–47^. Importantly, CD28 interaction with CD80/CD86 can be blocked by the CTLA4-Ig fusion protein abatacept, which is FDA approved for the treatment of rheumatoid arthritis, psoriatic arthritis, polyarticular juvenile idiopathic arthritis and acute graft versus host disease^48,49^. In myeloma, pre-clinical studies have shown that CTLA4-Ig in combination with melphalan can significantly reduce tumor burden^45^, leading to a phase II clinical trial of abatacept plus standard of care chemotherapy for treatment of patients with relapsed/refractory MM (NCT03457142).

Given the recently reported ∼3-fold improvement in overall response rate when abatacept was added to standard of care therapy^50^, we hypothesized that systemic blockade of CD28 would similarly sensitize MM to CAR T cell therapy. We reasoned that unlike endogenous T cells that require CD28 co-stimulation to mount an anti-tumor response, second generation CAR T cells receive co-stimulation directly from the CAR and would therefore be relatively unaffected by blockade of endogenous CD28. FDA approved CAR T cell products for MM, idacabtagene vicleucel and ciltacabtagene autoleucel, employ 4-1BB co-stimulatory domains, which have been shown to transduce a weaker signal than CD28 co-stimulatory domains in CD19 targeted CAR T cells^51,52^. In this context, the weaker 4-1BB co-stimulatory signal reduced CAR T cell exhaustion and enhanced *in vivo* persistence when compared to CD28 driven co-stimulation^53,54^. However, recent evidence suggests that enhanced CD28 signaling in CTLA-4 knockout 4-1BB co-stimulated (BBζ) CAR T cells improves their anti-tumor efficacy^55^. Moreover, endogenous tumor-reactive cytotoxic T cells require CD28 signaling to acquire effector properties in the tumor microenvironment^56^, suggesting that perhaps the CAR 4-1BB co-stimulatory domain alone is insufficient to stimulate potent anti-tumor activity.

In the current study, we employed human and mouse orthotopic models of multiple myeloma (MM) to directly test whether endogenous CD28 affected tumor control by BCMA targeted, 4-1BB co-stimulated CAR T cells. Somewhat unexpectedly, we found that continuous blockade of CD28 interaction with B7 proteins using abatacept significantly impaired CAR T cell control of MM growth, resulting in shorter survival of MM-bearing mice. Data indicate that abatacept primarily affected endogenous CD28 signaling on CAR T cells, as inducible deletion of CD28 from 4-1BB co-stimulated CAR T cells also reduced their *in vivo* anti-MM efficacy. Mechanistically, we provide evidence that endogenous CD28 signaling increases 4-1BB co-stimulated CAR T cell expansion in the MM bone marrow microenvironment (BME) by stimulating oxidative phosphorylation and maintaining redox balance. Transient inhibition of endogenous CD28 on 4-1BB co-stimulated CAR T cells resulted in decreased accumulation of CD4^+^ CAR T cells and release of inflammatory cytokines in the MM TME, without significantly impairing anti-MM activity. Collectively, our findings reveal that CAR T cell function is affected by endogenous CD28, which can potentially be transiently blocked to reduce toxic pro-inflammatory cytokine release while maintaining anti-tumor activity.

## Materials and Methods

### Cell lines

Parental 5TGM1 cells generously provided by G. David Roodman (Indiana University) were lentivirally transduced to express truncated human BCMA (hBCMA) and firefly luciferase to be used as target cells. Pure populations were achieved following fluorescence-activated cell sorting. MM.1S and U266 luciferase expressing clones were generated and supplied by Kelvin Lee (Indiana University). MM cell lines were maintained in RPMI 1640 (Gibco) supplemented with 10% heat-inactivated FBS (R&D Systems), 1% nonessential amino acids (Gibco), 1mM sodium pyruvate (Gibco), 10 mM HEPES (Gibco), 2 mM L-glutamine (Gibco) and 1% penicillin/streptomycin (Gibco). Cell lines were maintained in culture for 2-3 months at a time. 293T packaging cell lines were purchased from ATCC and maintained in DMEM, 10% FBS and 1% L-glutamine for 2-3 weeks prior to transient transfection. Stably expressing 293 Galv9 cells were kindly provided by Renier Brentjens. All cell lines were routinely tested for mycoplasma using the Lonza MycoAlert Detection Kit.

### Construct generation

Second generation CAR constructs were generously provided by Jonathan Bramson (McMaster University) and Renier Brentjens (Roswell Park) ^57^.

### Mouse αhBCMA CAR (hBCMAmBBmζ)

CAR encoding DNA was subcloned into the multiple cloning site of the pRV2011 retroviral vector which also contains an internal ribosome entry site (IRES) and Thy1.1. Murine CAR constructs consisted of an anti-human BCMA single chain variable fragment (scFv) (C11D5.3), CD8 hinge, CD28 transmembrane, 4-1BB signaling domain and CD3ζ activation domain as described previously ^58^.

### Mouse ahCD19 CAR (hCD19m28mζ)

Off-target CAR constructs consisted of an anti-human CD19 scFv (FMC63), CD28 transmembrane, CD28 signaling domain and CD3ζ activation domain.

### Human αhBCMA CAR (hBCMABBζ)

Antigen recognition was defined by an anti-human BCMA scFv previously reported. Human CAR constructs consisted of an CD8a hinge, CD8a transmembrane, CD28 *or* 4-1BB signaling domain and CD3ζ activation domain ^57,59^.

### hBCMA-tNGFR

Lentiviral expression plasmid, LeGO-Luc2, was a gift from Boris Fehse (Addgene plasmid #154006) which was further modified to co-express hBCMA-tNGFR. Murine 5TGM1 cells were transduced with lentiviral supernatant to drive expression of the target antigen, hBCMA, and permit *in vivo* imaging of tumor bearing animals. All constructs were verified by sanger sequencing.

### Mouse CAR T cell production

Mouse CAR T cell production was adapted from previous reports ^60^. Briefly, pan CD3^+^ murine T cells were isolated from single cell splenocyte suspensions of 6 – 12-week-old mice through a negative selection process (STEMCELL Technologies). T cells were activated with αCD3/αCD28 Dynabeads as specified by manufacturer’s instructions in the presence of 100 IU/mL recombinant mouse (rm) IL-2 and 10 ng/mL rmIL-7 (BioLegend) and cultured in RPMI1640 supplemented with 10% FBS, 2 mM L-glutamine, 10mM HEPES, 0.5% 2-mercaptoethanol, and 1% penicillin/streptomycin. Retroviral transduction was achieved by spinoculation of 3 × 10^6^ mouse T cells on retronectin-coated plates (Takara Bio) with neat retroviral supernatant harvested from 293T packaging cells (2000xg, 60 min., 30 °C) at 24 and 48 hr. post activation. CAR T cells were maintained at 1 × 10^6^ cells/mL for 7-10 days *in vitro* in the presence of rmIL-2 and rmIL-7 ^60^.

### Human CAR T cell production

Human CAR T cell production was adapted from previously published protocols ^57^. De-identified, healthy donor peripheral blood mononuclear cells (PBMCs) were obtained through the Roswell Park donor center under the approved protocol BDR 115919. PBMCs were isolated from whole blood through density gradient centrifugation. PBMCs were activated with T cell TransAct polymeric nanomatrix (Miltenyi Biotec) according to manufacturer’s specifications in the presence of 100 IU/mL recombinant human (rh) IL-2 and 10 ng/mL rhIL-7 (Peprotech). Spinoculation with neat 293 Galv9 retroviral supernatant was performed at 48, 72 and 96 hr. post activation (3200 rpm, 60 min., 30 °C). Human CAR T cells were expanded for 14 days and subsequently cryopreserved in 90% FBS, 10% DMSO.

### Cytotoxicity assays

*In vitro* CAR T cell killing assays were performed using firefly luciferase expressing target cells. Briefly, 2 × 10^4^ target cells were seeded in 96 well plates, varying numbers of CAR T cells were added to assess CAR T cell mediated killing within the linear range. Target cell viability following co-culture incubation was determined using the ONE-glo luciferase reporter assay (Promega). For cytokine secretion assays, supernatants were collected 24 hrs. after co-culture and analyzed on a Luminex xMAP INTELLIFLEX system.

### Flow cytometry

Data was acquired on either a LSR Fortessa (BD Biosciences) or Cytek Aurora full spectrum analyzer (Cytek Biosciences). Analysis was performed using FlowJo (Tree Star Inc.) or FCS Express software (De novo Software). Briefly, cell suspensions were harvested, washed, and stained with fixable live/dead blue (Invitrogen) in PBS followed by surface antibody staining in FACS buffer (1% BSA, 0.1% sodium azide in PBS). Antibodies were titrated for optimal staining for 20 min. at 4°C. Intracellular cytokine staining was conducted following fixation and permeabilization according to manufacturer’s instructions (BioLegend). Antibodies used in phenotypic analysis are included in Supplemental Table 1.

### Cytokine analyses (Luminex and Isoplexis)

Co-culture supernatants were immediately snap frozen and stored at −80 °C until Luminex assays were run. Detection of mouse CAR T cell cytokine production was performed using the MILLIPLEX Th17 premixed panel and acquired on a Luminex xMAP INTELLIFLEX system. Data was analyzed using the Belysa^®^ Immunoassay Curve Fitting Software (Millipore Sigma). Cytokine production within the BM TME was evaluated using Isoplexis’ CodePlex secretome chips. Briefly, bilateral, tumor-bearing hind limbs (femur and tibia) were harvested, and BM was collected into 30 µL PBS. Soluble phase fractions were stored at −80 °C in low-bind Eppendorf tubes until loaded onto a CodePlex secretome chip and analyzed in an IsoLight instrument.

### Seahorse

The day prior to assay CAR T cells were stimulated overnight at an E:T ratio of 2:1. CelllZlTaklZlCoated XF96 microplate was prepared to support testing of cells grown in suspension. Sensor cartridge was hydrated in XF Calibrant and incubated overnight at 37 °C in a non-CO_2_ incubator.

On the day of the experiment Seahorse XF DMEM Medium pH 7.4 (Agilent Technologies) was supplemented with 10 mM glucose, 1 mM pyruvate and 2 mM glutamine for oxygen consumption rate (OCR) examination and 2mM glutamine for extracellular acidification rate (ECAR) examination and pre-warmed to 37 °C. Suspension cells were harvested, washed in 1X PBS, resuspended in the prepared assay media, and gently seeded on the CelllZlTaklZlCoated plate at 2 × 10^5^ cells/well. The seeded plate was incubated in a 37 °C non-CO_2_ incubator for 1 hour prior to the assay. During the incubation, test compounds specific to the assay type were prepared and added to the ports of the hydrated cartridge. The loaded cartridge was moved to the XFe96 Analyzer and initial calibration was performed. Following the 1-hour incubation the Cell-Tak plate was transferred to the Xfe96 Analyzer and the assay was initiated according to manufacturer’s recommendations.

All concentrations shown represent final well concentrations:

### Mito Stress Test (OCR examination)

Oligomycin (2.0 µM), FCCP (1.0 µM) and Rotenone/Antimycin A (0.5 µM)

### Glycolysis Stress Test (ECAR examination)

Glucose (10 mM), Oligomycin (1.0 µM) and 2DG (50 mM)

### T Cell Metabolic Fitness Test

First injection of this assay included substrate pathway specific inhibitors:

Etomoxir (4.0 µM), long chain fatty acid oxidation

UK5099 (2.0 µM), glucose/pyruvate oxidation

BPTES (3.0 µM) glutamine oxidation

For controls, assay medium was used in port A instead of inhibitors.

Following inhibitor injections Oligomycin (1.5 µM), BAM15 (2.5 µM) and Rotenone/Antimycin A (0.5 µM) were added to all wells.

The last injection of each assay included Hoechst 33342 Nuclear Stain to facilitate Normalization via fluorescent imaging and cell counting supported by the BioTek Cytation 5 Cell Imaging Multimode Reader. Data was analyzed using the Wave 2.6.1 software and the Seahorse Analytics cloud-based resource.

### qRT-PCR

RNA was isolated using the miRNeasy Mini Kit (Qiagen) and cDNA was synthesized using SuperScript IV Reverse Transcriptase. Contaminating DNA was removed using Rnase-Free Dnase (Qiagen) and qPCR was performed using the QuantStudio 6 Flex Real-Time PCR System with SYBR green (ThermoFisher Scientific). Expression was normalized to TBP, and relative expression was calculated using the ΔΔCT formula. Primer sequences are listed in Supplemental Table 2.

### NAD^+^/NADH Quantitation

NADH:NAD^+^ ratios were determined using the NAD/NADH-Glo assay (Promega) according to manufacturer’s instructions. Briefly, 1 × 10^5^ stimulated CAR T cells were washed with 1x PBS prior to cell lysis. NAD^+^ and NADH levels were quantified independently using acid/base treatment. Luminescence values were read on a BioTek Synergy H1 plate reader (Agilent).

### Mice and in vivo models

#### Systemic Tumor Models

NOD scid gamma (NSG) mice (NOD.Cg*Prkdc^scid^ Il2rg^tm1Wjl^*/SzJ) mice ages 6-12 weeks were purchased from the Comparative Oncology Shared Resource in-house mouse colony at Roswell Park. RAG2^-/-^ (B6.Cg-*Rag2^tm^*^1^*^.1Cgn^*/J) mice were purchased from the Jackson Laboratory and subsequently bred in our facility under the approved protocol 1425M. NSG mice were intravenously injected with 1 × 10^6^ MM.1S-Luc at week −4 and 3 × 10^6^ CAR T cells at week 0. RAG2^-/-^ mice were injected with 2 × 10^6^ 5TGM1^hBCMA^-Luc at week −2 to and 3 × 10^6^ CAR T cells at week 0 to compensate for differences in tumor engraftment rate amongst the two models. Bioluminescence was measured 2x/week using an IVIS^®^ Spectrum In Vivo Imaging System (PerkinElmer) to assess tumor burden. Mice were injected with 150 mg Luciferin/kg of body weight and briefly anesthetized through isoflurane inhalation during image acquisition. Data was analyzed on the Living Image analysis software (PerkinElmer). In some settings, retroorbital blood collection was performed 1 week after CAR T cell infusion to examine CAR T cell frequency in circulation. Mice were monitored daily for signs of deteriorating condition or disease progression including decreased activity, hunched posture, ruffled coat, or hind limb paralysis and euthanized upon veterinary recommendation.

All animal studies were performed in accordance with the Roswell Park Comprehensive Cancer Center Institutional Animal Care and Use Committee guidelines under the approved protocol 1094M.

#### CD28^iKO^

CD28^iKO^ mice were generated by Ozgene (Australia). LoxP sites flanking exon 2 and 3 of the CD28 gene were introduced to allow for Cre-mediated deletion of the CD28 gene. Mice were generously provided by Kelvin Lee (Indiana University) and subsequently bred in-house under protocol 1425M. Splenocytes were isolated as previously described and CAR T cells were expanded in the presence of 250 nM 4-hydroxytamoxifen (Sigma-Aldrich) for 4 days to induce CD28 deletion.

#### T-lux

Transgenic T-lux mice generated by Casey Weaver at the University of Alabama at Birmingham were acquired by Robert McGray (Roswell Park) under a material transfer agreement (MTA). Mice were utilized as splenocyte donors for CAR T cell manufacturing for *in vivo* imaging of CAR T cell trafficking and expansion.

#### Statistical analyses

All statistical analyses were performed using GraphPad Prism software. Data points represent independent biological replicates. Error bars represent standard deviation unless otherwise stated. Statistical significance between groups was determined by paired or unpaired Student’s t test, one-way or two-way ANOVA. Survival analysis was performed using a log-rank (Mantel-Cox) test. A p value ≤0.05 is considered significant: *p<0.05, **p<0.01, ***p<0.001, ****p<0.0001.

## Results

### Blockade of endogenous CD28 impairs hBCMABB***ζ*** CAR T cell anti-MM activity

Since inhibition of the CD28 survival signal in multiple myeloma (MM) cells sensitizes them to chemotherapy^45,46^, we sought to determine whether CD28 inhibition similarly sensitized MM cells to killing by CAR T cells. Human CAR T cells targeting BCMA and containing a 4-1BB/CD3ζ intracellular signaling domain (hBCMABBζ) similar to FDA approved CAR products for MM (Fig. 1A) were generated from healthy donor peripheral blood mononuclear cells transduced with a previously described retroviral vector ^57^ (Supplemental Fig. 1A – 1E). Co-culture of hBCMABBζ CAR T cells with human MM cell lines MM.1S or U266, which differ in their expression profiles of CD28 and B7 ligands (MM.1S = CD28^+^, CD86^+^, CD80^-^ & U266 = CD28^+^, CD86^-^,CD80^-^; Supplemental Fig. 1F), resulted in cytotoxicity across a range of effector to target ratios (Fig. 1B). Intriguingly, addition of abatacept to co-cultures mildly enhanced sensitivity of CD86^+^ MM.1S, but not CD86^-^ U266, to hBCMABBζ CAR T cell killing, indicating that blocking CD28-CD86 interactions on MM cells may sensitize them to CAR T cell therapy (Fig. 1B). In agreement with potential MM sensitization to CAR T killing, abatacept did not alter hBCMABBζ CAR T cell production of effector cytokines or proinflammatory molecules including interferon-gamma (IFN-γ), tumor necrosis factor alpha (TNFα) or granulocyte-macrophage colony-stimulating factor (GM-CSF) in co-culture assays (Supplemental Fig. 1G).

**Figure 1:**
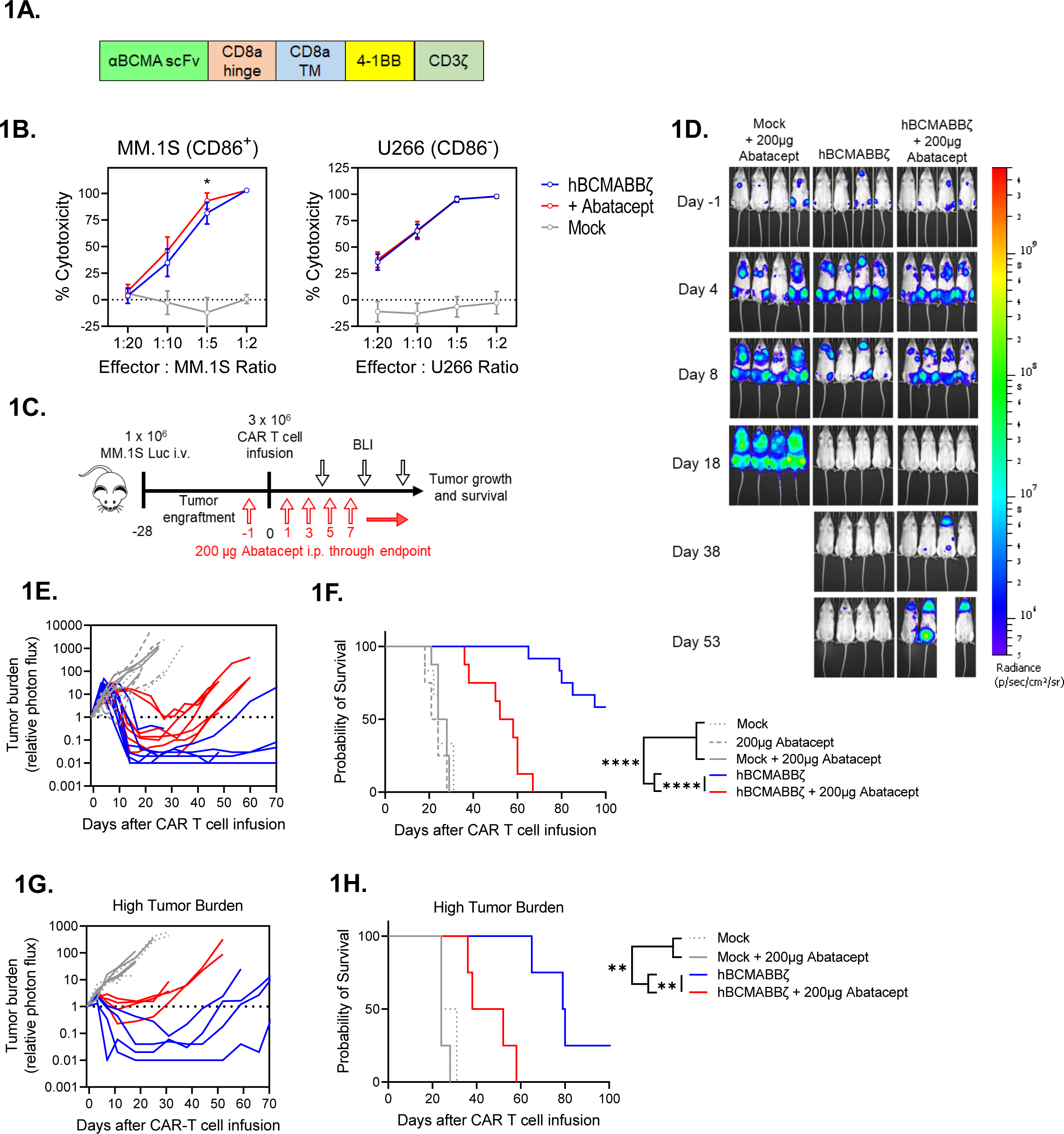
Endogenous CD28 blockade impairs hBCMABBζ CAR T cell anti-myeloma efficacy. **(A)** Schematic of second-generation retroviral CAR construct used to generate BCMA targeted human CAR T cells. **(B)** CAR T cell cytotoxic activity during a 24-hr. co-culture with luciferase-tagged MM.1S (left) or U266 (right) myeloma cells ± abatacept. Data are shown as mean ± SD and representative of hBCMABBζ CAR T cells generated from 3 healthy donors. *p<0.05 by two-way analysis of variance (ANOVA) with Tukey’s multiple comparison test. **(C)** Diagram of experimental setup used to evaluate hBCMABBζ CAR T + abatacept therapy in a human MM xenograft model. NSG mice were intravenously inoculated with 1 × 10^6^ MM.1S-luc myeloma cells on day −28 and treated with CAR T cells on day 0. Mice received 200 µg abatacept 3x/week beginning the day before infusion and continuing through endpoint. Tumor burden was monitored by IVIS bioluminescent imaging (BLI) 2x/week. **(D)** Representative bioluminescent images of MM.1S bearing mice on specified days following CAR T cell infusion. **(E)** Tumor burden expressed as relative photon flux measured by BLI from MM.1S-luc bearing mice treated with hBCMABBζ CAR T or control T cells ± abatacept (200 µg, 3x/week). Each line represents an individual mouse (n = 7 mice per CAR T cell treated group). **(F)** Kaplan – Meier analysis of survival of hBCMABBζ CAR T or control T cells ± abatacept (200 µg, 3x/week) treated MM.1S-luc bearing mice (n = 8 - 12 mice per CAR T cell treated group). Median survival of hBCMABBζ CAR T treated mice was >100 days post CAR T cell infusion vs. 55 days for hBCMABBζ CAR T + abatacept treated mice. ****p<0.0001 by log-rank Mantel-Cox test. **(G)** Tumor burden expressed as relative photon flux measured by BLI from MM.1S high tumor burden mice (inoculated on day −35) treated with hBCMABBζ CAR T or control T cells ± abatacept (200 µg, 3x/week). Each line represents an individual mouse (n = 4 mice per CAR T cell treated group). **(H)** Kaplan – Meier analysis of survival of hBCMABBζ CAR T or control T cells ± abatacept (200 µg, 3x/week) treated MM.1S-luc high tumor burden mice (n = 4 mice per group). Median survival of hBCMABBζ CAR T treated mice was 80 days vs. 45 days for hBCMABBζ CAR T + abatacept treated mice. ***p<0.0001 by log-rank Mantel-Cox test.

Due to the modest capacity of abatacept to enhance CAR T cell-mediated cytotoxicity *in vitro*, we evaluated the ability of abatacept to enhance hBCMABBζ CAR T cell control of orthotopic CD28^+^, CD86^+^ myeloma. Luciferase tagged MM.1S (MM.1S-Luc) cells were implanted i.v. into NSG hosts followed four weeks later by infusion of 3 × 10^6^ hBCMABBζ CAR T cells ± 3x/weekly injections of abatacept continued until endpoint (Fig. 1C). Bioluminescence imaging was used to confirm bone marrow engraftment and to normalize average tumor burden across groups immediately prior to therapy. Following hBCMABBζ CAR T infusion, MM.1S burden was assessed by serial bioluminescence imaging. MM regression was observed in all hBCMABBζ CAR T cell treated mice, with most mice apparently tumor free 2 to 3 weeks following infusion (Fig. 1D). Unexpectedly, MM relapse was more rapidly seen in mice receiving abatacept + hBCMABBζ CAR T cells compared to those receiving single agent hBCMABBζ CAR T cells (Fig. 1E), resulting in significantly shorter survival of MM.1S bearing mice in the abatacept + hBCMABBζ CAR T group (Fig. 1F).

Prior work has demonstrated that CD28 can contribute to an immunosuppressive MM BME by interacting with CD80/CD86 on bone marrow resident DCs and inducing production of IL-6 and the tryptophan metabolizing enzyme, indoleamine 2,3-dioxygenase (IDO)^44^. In the low tumor burden setting, abatacept may function through ligation of B7 family proteins on BMDCs to create an immunosuppressive MM BME. We therefore repeated CAR T cell ± abatacept treatment regimen in a high MM.1S tumor burden setting in which an immunosuppressive MM BME should already be established. We found that abatacept similarly accelerated relapse following hBCMABBζ CAR T cell infusion in the high tumor burden setting (Fig. 1G) and shortened overall survival (Fig. 1H) suggesting that induction of immunosuppression in the MM bone marrow microenvironment was likely not the primary effect of abatacept exposure.

### Endogenous CD28 enhances 4-1BB co-stimulated CAR T cell efficacy

Despite the very clear reduction to *in vivo* CAR T cell efficacy imparted by continuous abatacept treatment, data shown in Fig. 1 does not differentiate effects of abatacept on cells in the MM BME versus effects of blocking endogenous CD28 on CAR T cells. To examine CAR T cell intrinsic effects of endogenous CD28, we generated a tamoxifen inducible CD28 knockout mouse model (CD28^iKO^) by crossing CD28-floxed mice to mice expressing CreERT2 from the ROSA26 locus^61^. Following hBCMAmBBmζ CAR transduction of CD28^iKO^ or littermate control T cells lacking CreERT2 expression, 4-hydroxytamoxifen was introduced into culture media for 4 days to induce CD28 deletion (Fig. 2A). Surface protein expression was evaluated by flow cytometry over the course of CAR T cell expansion and immediately prior to functional assessment. Importantly, CD28 surface expression was reduced to near background levels in CD28^iKO^ CAR T cells by 4-OHT exposure (Fig. 2B,2C), while CD4: CD8 ratio and CAR expression was unaffected (Supplemental Fig. 2A – 2C). To interrogate functionality of murine CAR T cells we engineered syngeneic 5TGM1 MM cells to express a chimeric hBCMA-tNGFR target antigen (5TGM1^hBCMA^) ^62–65^ (Supplemental Fig. 2D, 2E). Coupling the extracellular domains of hBCMA to a signaling deficient NGFR transmembrane domain allowed us to uncouple the target function of BCMA from its survival signal. In co-culture assays, CD28^iKO^ CAR T cells were nearly as effective as control CD28^fl/fl^ CAR T cells at killing 5TGM1^hBCMA^ myeloma cells, indicating that endogenous CD28 does not directly impact CAR T cell cytotoxicity (Fig. 2D). CD28^fl/fl^ and CD28^iKO^ CAR T cells also produced comparable amounts of proinflammatory cytokines when stimulated by 5TGM1^hBCMA^ *in vitro* (Fig. 2E).

**Figure 2:**
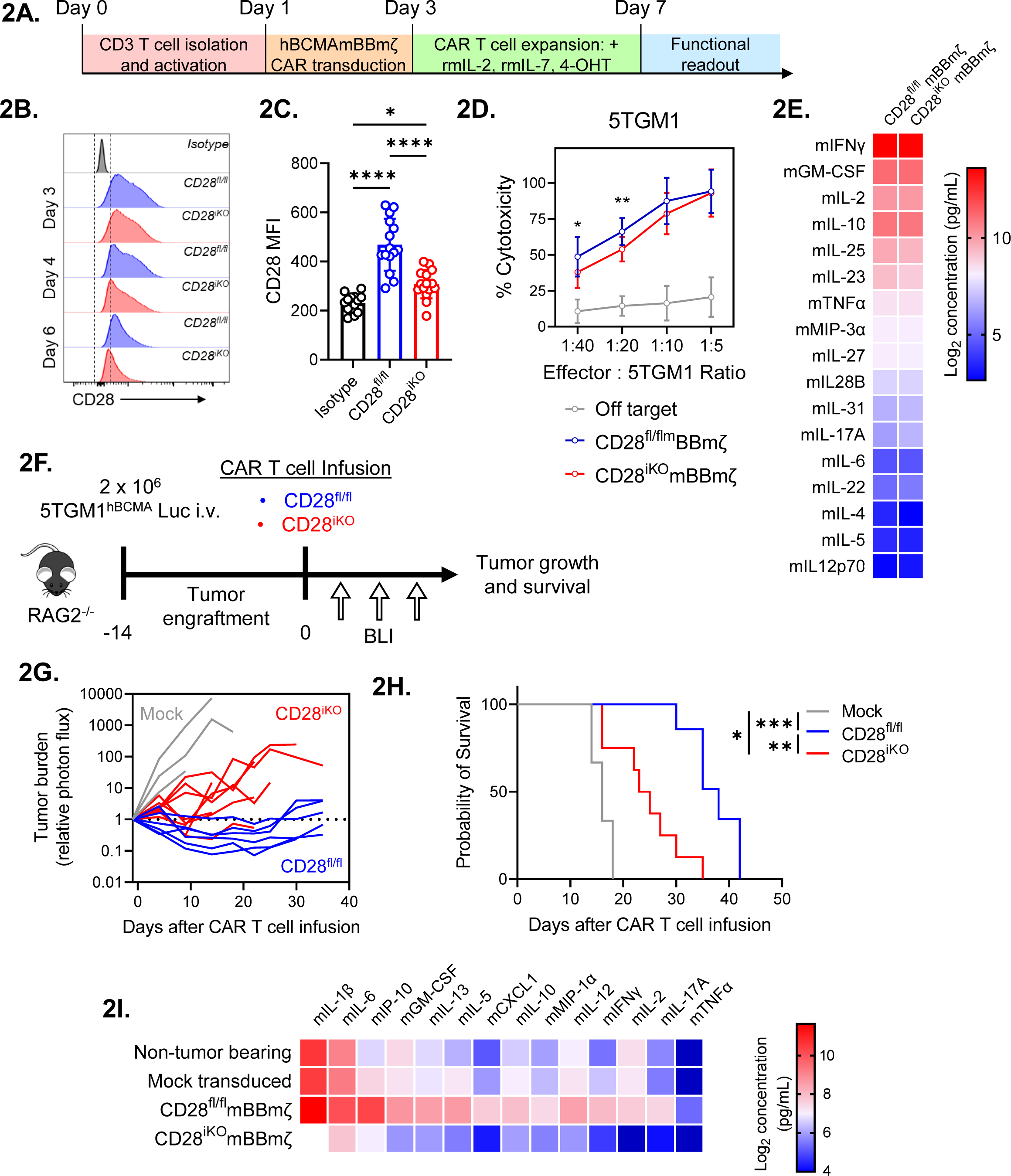
CD28^iKO^ hBCMAmBBmζ CAR T cells are functionally impaired *in vivo*. **(A)** Schematic depicting the process of manufacturing mouse CD28 knockout (CD28^iKO^) hBCMAmBBmζ CAR T cell. **(B)** Surface CD28 protein expression on CD28^fl/fl^ versus CD28^iKO^ mouse T cells during CAR T cell manufacture. **(C)** Median fluorescent intensity (MFI) of CD28 measured by flow cytometry at the conclusion of CD28^fl/fl^ versus CD28^iKO^ mouse CAR T manufacture. Data shown as mean ± SD from 10+ independent experiments. *p<0.05, ****p<0.0001 by one-way ANOVA. **(D)** CD28^fl/fl^ versus CD28^iKO^ hBCMAmBBmζ CAR T cell cytotoxic activity during 24 hr. co-culture with luciferase-tagged 5TGM1^hBCMA^ mouse myeloma cells. Cell viability was assessed by luciferase assay. Data are shown as mean ± SD and representative of at least 3 independent experiments. *p<0.05, **p<0.01 by two-way ANOVA with Tukey’s multiple comparison test. **(E)** Heatmap representation of culture supernatant mouse cytokine concentrations measured by multiplexed Luminex assays at the conclusion of a 24-hr. co-culture of hBCMAmBBmζ CD28^fl/fl^ or CD28^iKO^ CAR T cells with 5TGM1^hBCMA^ myeloma cells. Log_2_ transformed cytokine concentrations represent the mean of 4 independent experiments. **(F)** Diagram of experimental setup used to evaluate CD28^fl/fl^ versus CD28^iKO^ hBCMAmBBmζ CAR T cell therapy in a mouse MM xenograft model. RAG2^-/-^ mice were inoculated intravenously with 2 × 10^6^ 5TGM1^hBCMA^-luc cells on day −14 and treated with CD28^fl/fl^ or CD28^iKO^ CAR T cells on day 0. Tumor burden was monitored by IVIS bioluminescent imaging (BLI) 2x/week through endpoint. **(G)** Tumor burden expressed as relative photon flux measured by BLI from 5TGM1^hBCMA^-luc bearing mice treated with CD28^fl/fl^ or CD28^iKO^ hBCMAmBBmζ CAR T or control T cells. Each line represents an individual mouse (n = 6 mice per CAR T cell treated group). **(H)** Kaplan – Meier analysis of survival of CD28^fl/fl^ or CD28^iKO^ hBCMAmBBmζ CAR T cell or control T cell treated 5TGM1^hBCMA^-luc bearing mice (n = 6 mice per CAR T cell treated group). Median survival of CD28^fl/fl^ hBCMAmBBmζ CAR T treated mice was 38 days post-CAR T cell infusion vs. 24 days for CD28^iKO^ hBCMAmBBmζ CAR T treated mice. *p<0.05, **p<0.01, ***p<0.001 by log-rank Mantel-Cox test. **(I)** Heatmap representation of cytokine levels in the MM BME 7 days following infusion of CD28^fl/fl^ or CD28^iKO^ hBCMAmBBmζ CAR T cells into 5TGM1^hBCMA^-luc bearing mice. Bilateral hind limbs were harvested, and BM was flushed into 15 µL PBS for multiplexed cytokine analysis. Log_2_ transformed cytokine concentrations represent the mean of 3 mice per group.

In contrast to *in vitro* findings, CD28^iKO^ hBCMAmBBmζ CAR T cells differed greatly from hBCMAmBBmζ CAR T cells generated from littermate controls in their ability to control *in vivo* myeloma growth. CD28^iKO^ hBCMAmBBmζ CAR T cells transiently controlled systemic growth of luciferase labeled 5TGM1^hBCMA^ myeloma in RAG2^-/-^ mice while CD28^fl/fl^ littermate control hBCMAmBBmζ CAR T cells demonstrated extended myeloma control (Fig. 2F, 2G). As a result, the median survival of 5TGM1^hBCMA^ myeloma bearing mice treated with CD28^fl/fl^ hBCMAmBBmζ CAR T cells was nearly twice as long as those treated with CD28^iKO^ hBCMAmBBmζ CAR T cells (Fig. 2G, 2H), mirroring effects of abatacept blockade of endogenous CD28 signaling (Fig. 1E, 1F). Moreover, pro-inflammatory cytokines in the MM BME of CD28^iKO^ hBCMAmBBmζ CAR T cell treated mice were substantially reduced (Fig. 2I). These data indicate that CD28^iKO^ CAR T cells did not induce a proinflammatory MM BME despite being able to readily produce pro-inflammatory cytokines in response to CAR ligation (Fig. 2E).

### Endogenous CD28 supports CAR T cell oxidative metabolism

CD28 controls metabolic reprogramming of activated T cells to enhance production of pro-inflammatory cytokines and anti-tumor immunity^66–69^. Since CAR T cell efficacy is linked to the metabolic state of infused cells^70^, we evaluated glycolytic and mitochondrial metabolism of unstimulated and 5TGM1^hBCMA^ stimulated CD28^fl/fl^ and CD28^iKO^ hBCMAmBBmζ CAR T cells using Seahorse assays. Changes in extracellular acidification rate (ECAR) in response to glucose addition or inhibition of mitochondrial ATP synthesis were equivalent between CD28^fl/fl^ and CD28^iKO^ CAR T cells (Fig. 3A, 3B), indicating that mBBmζ CAR signaling was sufficient to induce glycolytic metabolism. In contrast, CD28^iKO^ hBCMAmBBmζ CAR T cells displayed reduced oxygen consumption rate (OCR) and this pattern differed based on stimulation (Fig. 3C,3D). Basal and uncoupled OCR were decreased in unstimulated CD28^iKO^ CAR T cells while uncoupled OCR and spare respiratory capacity (SRC) were decreased in stimulated CD28^iKO^ CAR T cells. Reduced OCR in CD28^iKO^ CAR T cells is consistent with the established role of CD28 in priming mitochondria to support a robust recall response in of memory CD8 T cells^67^. Yet in contrast to memory CD8 T cells, reduced mitochondrial OCR in CD28^iKO^ CAR T cells did not result from diminished fatty acid oxidation nor from an inability of CD28^iKO^ CAR T cells to oxidize other major anapleurotic substrates glucose and glutamine (Supplemental Fig. 3A). Moreover, no difference in mitochondria content was observed when comparing CD28^iKO^ and CD28^fl/fl^ hBCMAmBBmζ CAR T cells (Supplemental Fig. 3B), further suggesting that endogenous CD28 signaling regulates mitochondrial oxidative phosphorylation in 4-1BB co-stimulated CAR T cells.

**Figure 3:**
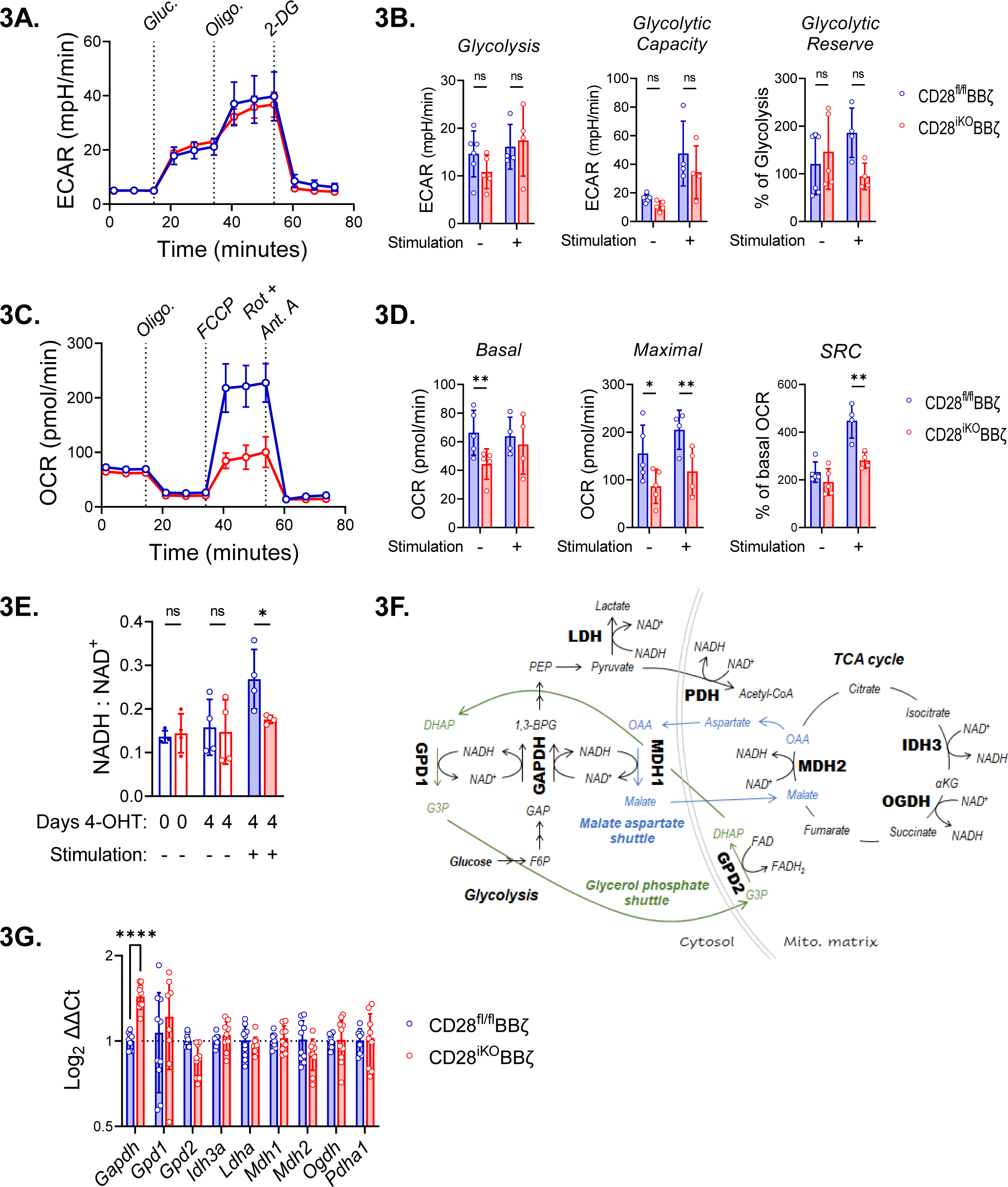
Perturbation of oxidative metabolism and redox homeostasis in stimulated CD28^iKO^ hBCMAmBBmζ CAR T cells. **(A)** Representative Seahorse Glycolysis Stress Test performed 24 hr. after stimulation of CD28^fl/fl^ or CD28^iKO^ hBCMAmBBmζ CAR T cells with 5TGM1^hBCMA^ myeloma cells. Connected data points represent mean extracellular acidification rate (ECAR) ± SD of four technical replicates at indicated time points during a representative experiment that was repeated at least 3 times. Dashes indicate the timing of glucose, oligomycin (Oligo) and 2-deoxyglucose (2-DG) injection. **(B)** Quantified rates of glycolysis (left), glycolytic capacity (middle), and glycolytic reserve (left) in CD28^fl/fl^ or CD28^iKO^ hBCMAmBBmζ CAR T cells ± 24-hr. stimulation with 5TGM1^hBCMA^ myeloma cells. Bars represent mean ± SD, dots represent independent experiments. ns = not significant by one-way ANOVA. **(C)** Representative Seahorse Mito Stress Test performed 24 hr. after stimulation of CD28^fl/fl^ or CD28^iKO^ hBCMAmBBmζ CAR T cells with 5TGM1^hBCMA^ myeloma cells. Connected data points represent mean oxygen consumption rate (OCR) ± SD of four technical replicates at indicated time points during a representative experiment that was repeated at least 3 times. Dashes indicate the timing of oligomycin (Oligo), FCCP, and rotenone (Rot) + antimycin A (Ant. A) injection. **(D)** Quantified basal OCR (right), uncoupled maximal respiration (middle), and spare respiratory capacity (SRC) expressed as percent of basal OCR (left) in CD28^fl/fl^ or CD28^iKO^ hBCMAmBBmζ CAR T cells ± 24 hr. stimulation with 5TGM1^hBCMA^ myeloma cells. Bars represent mean ± SD, dots represent independent experiments. *p<0.05, **p<0.01 by one-way ANOVA. **(E)** Ratio of NADH to NAD^+^ ratios in CD28^fl/fl^ or CD28^iKO^ hBCMAmBBmζ CAR T cells prior to 4-OHT mediated CD28 deletion or ± 24 hr. stimulation with 5TGM1^hBCMA^ myeloma cells. Bars represent mean ± SD from 3 independent experiments. *p<0.05 by paired Student’s t test. **(F)** Diagrams depicting enzymes of central carbon metabolism that interconvert NADH and NAD^+^. **(G)** Relative expression of mRNAs coding enzymes depicted in (F) in CD28^iKO^ versus CD28^fl/fl^ hBCMAmBBmζ CAR T cells following 24 hr. stimulation with 5TGM1^hBCMA^ myeloma cells. Bars represent mean log_2_ transformed ΔΔCt values ± SD, dots technical replicates pooled from 3 independent experiments. *Tbp* and *Actb* were used as endogenous controls. ****p<0.0001 by two-way ANOVA.

Mitochondrial oxidative phosphorylation relies on the electron carriers NADH to donate electrons to complex I and FADH2 to donate electrons to complex II of the electron transport chain (ETC), driving mitochondrial oxygen consumption and creating a proton gradient across the inner mitochondria membrane to fuel ATP synthase. Deletion of endogenous CD28 did not alter the contribution of complex I nor complex II to mitochondrial oxygen consumption by hBCMAmBBmζ CAR T cells (Supplemental Fig. 3C). Interestingly however, the ratio of NADH to NAD^+^ was increased in target cell stimulated CD28^fl/fl^ when compared to CD28^iKO^ hBCMAmBBmζ CAR T cells (Fig. 3E), indicating that endogenous CD28 signaling increases ETC substrate availability in 4-1BB co-stimulated CAR T cells. Several crucial metabolic enzymes reduce NAD^+^ to NADH or oxidize NADH to NAD^+^ (Fig. 3F), thereby maintaining redox balance. Quantitative RT-PCR revealed that among these enzymes, only *Gapdh* gene expression was altered in CD28^iKO^ hBCMAmBBmζ CAR T cells (Fig. 3g). Expression of the NADP^+^ reducing enzymes *Idh1* and *Idh2* were also slightly increased in CD28^iKO^ hBCMAmBBmζ CAR T cells (Supplemental Fig. 3D). Differences in gene expression, if reflected in functional enzyme changes, occur in the opposite direction of what would be expected based on differences in NADH to NAD^+^ ratio between CD28^fl/fl^ and CD28^iKO^ CAR T cells, indicating that gene expression changes are unlikely to explain the difference in redox state when CD28 is knocked out of hBCMAmBBmζ CAR T cells.

### Endogenous CD28 enhances CAR T cell expansion in the MM BME

Due to the known influence of mitochondrial respiration and redox balance on T cell proliferation^71,72^, we evaluated proliferation of CD28^fl/fl^ and CD28^iKO^ CAR T cells. No difference in proliferation of CD28^fl/fl^ and CD28^iKO^ CAR T cells was observed over the course of *ex vivo* CAR manufacture (Fig. 4A). Similarly, expression of the proliferation marker Ki67 induced by co-culture of CD28^fl/fl^ or CD28^iKO^ hBCMAmBBmζ CAR T cells with 5TGM1^hBCMA^ target cells was similar (Fig. 4B). However, when hBCMAmBBmζ CAR T cells in the MM BME or peripheral blood were enumerated 7 days after adoptive transfer into 5TGM1^hBCMA^ bearing mice (Fig. 4C), a large decrease in CD4^+^ CD28^iKO^ CAR T cells was observed (Fig. 4D, 4E, Supplemental Fig. 4A). Importantly, CD28^iKO^ hBCMAmBBmζ CAR T cells maintained low/negative CD28 surface expression in the MM BME (Fig. 4F). Abatacept similarly reduced the frequency of human CD4^+^ hBCMABBζ CAR T cells in the MM BME of MM.1S bearing mice (Fig. 4G), while it had no effect on human CAR T cells in the peripheral blood (Fig. 4H). The frequency of CD8^+^ CAR T cells in the MM BME or peripheral blood was unaffected by CD28 knockout or abatacept treatment (Fig. 4D – 4H).

**Figure 4:**
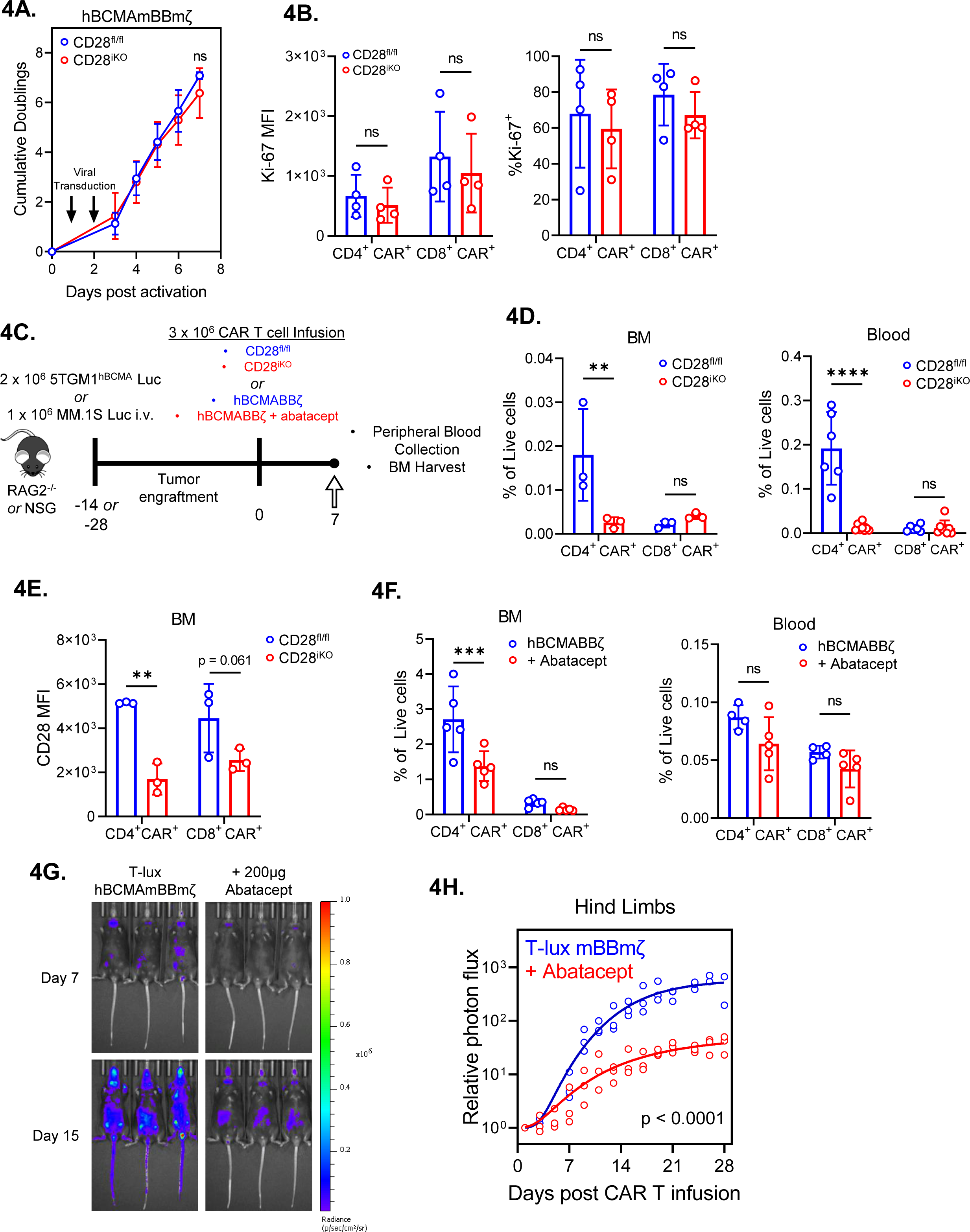
Diminished *in vivo* expansion of 4-BB co-stimulated CD28^iKO^ CAR T cells. **(A)** Expansion of CD28^fl/fl^ versus CD28^iKO^ hBMCAmBBmζ CAR T cells over the course of manufacturing. ns = not significant by two-way ANOVA. **(B)** Expression of the proliferation marker Ki-67 in CD28^fl/fl^ versus CD28^iKO^ hBCMAmBBmζ CAR T cells following 24 hr. stimulation with 5TGM1 target cells as assessed by flow cytometric analysis. ns = not significant by one-way ANOVA. **(C)** Diagram of experimental setup. 5TGM1^hBCMA^ bearing RAG2^-/-^ mice were treated with CD28^fl/fl^ or CD28^iKO^ hBCMAmBBmζ CAR T cells (4D and 4E) or MM.1S bearing NSG mice were treated with hBCMABBζ CAR T cells ± 200μg abatacept on days −1, 1, 3, 5, 7 (4F) and euthanized 7 days post adoptive transfer for blood collection and hind limb BM harvest. **(D)** CAR T cell frequency assessed by flow cytometry in bone marrow (BM, left) and peripheral blood (right) one week after adoptive transfer of CD28^fl/fl^ or CD28^iKO^ hBCMAmBBmζ CAR T cells into 5TGM1^hBCMA^ myeloma bearing mice. Bars represent mean ± SD, dots indicate individual mice (n = 3-6 mice per group). **p<0.01, ****p<0.0001 by two-way ANOVA with Tukey’s multiple comparison test. **(E)** CD28 surface protein expression on BM-infiltrating CD28^fl/fl^ or CD28^iKO^ hBCMAmBBmζ CAR T cells one week after adoptive transfer into 5TGM1^hBCMA^ myeloma bearing mice. **(F)** CAR T cell frequency assessed by flow cytometry in bone marrow (BM, left) and peripheral blood (right) one week after adoptive transfer of hBCMABBζ CAR T cells into MM.1S myeloma bearing mice ± abatacept. CAR T population identified by surface staining and analyzed by flow cytometry. Bars represent mean ± SD, dots indicate individual mice (n = 4-5 mice per group). ***p<0.001 by two-way ANOVA with Tukey’s multiple comparison test. **(G)** Representative IVIS bioluminescence images of T-lux luciferase expressing hBCMAmBBmζ CAR T cells ± abatacept (200μg, 3x/week) on day 7 or day 15 after infusion. **(H)** Quantification of photon flux by IVIS imaging within the hind limb region of interest (ROI) over a four-week period following T-lux hBCMAmBBmζ CAR T cell infusion ± abatacept (200μg, 3x/week) into 5TGM1^hBCMA^ myeloma bearing mice. Dots represent individual mice (n = 3 mice per group), lines are best-fit sigmoidal curves, and significance was determined by mixed effects modeling.

The observed reduction in CD4^+^ CAR T cells in the MM BME upon deletion or blockade of endogenous CD28 led us to test whether endogenous CD28 signaling contributes to *in vivo* expansion of luciferase expressing hBCMABBζ CAR T cells generated from T-lux mice^73^. Approximately one week after infusion, which aligns with the kinetics of tumor regression, T-lux hBCMAmBBmζ CAR T cell luminescence within the hind limbs of 5TGM1^hBCMA^ myeloma bearing mice rapidly increased with a signal plateau observed approximately 1 week later (Fig. 4I, 4J). Abatacept treatment significantly blunted *in vivo* expansion of T-lux hBCMAmBBmζ CAR T cells in the MM BME (Fig. 4I), suggesting that CD28 signaling supports *in vivo* expansion of 4-1BB co-stimulated CAR T cells.

### Transient CD28 blockade reduces inflammatory cytokines in the MM BME

Since endogenous CD28 promoted *in vivo* hBCMAmBBmζ CAR T cell expansion and inflammatory cytokine production in the MM BME, we sought to test whether abatacept blockade of CD28 ligation could lessen the severity of CAR T associated cytokine release. To this end, myeloma bearing mice were treated with abatacept for 1 week following CAR T cell infusion (Fig. 5A). At this early timepoint, abatacept had no effect on anti-tumor activity of BCMA targeted human or mouse CAR T cells (Fig. 5B, Supplemental Fig. 4B) and only a very minor effect on MM BME levels of human inflammatory cytokines in the MM BME (Fig. 5C). Since human cytokines could only come from CAR T cells or MM.1S cells, and most cytokines measured are not known to be made by MM cells, we concluded that CD28 blockade with abatacept did not affect *in vivo* CAR T cell cytokine secretion nor anti-tumor activity in the first week following infusion.

**Figure 5:**
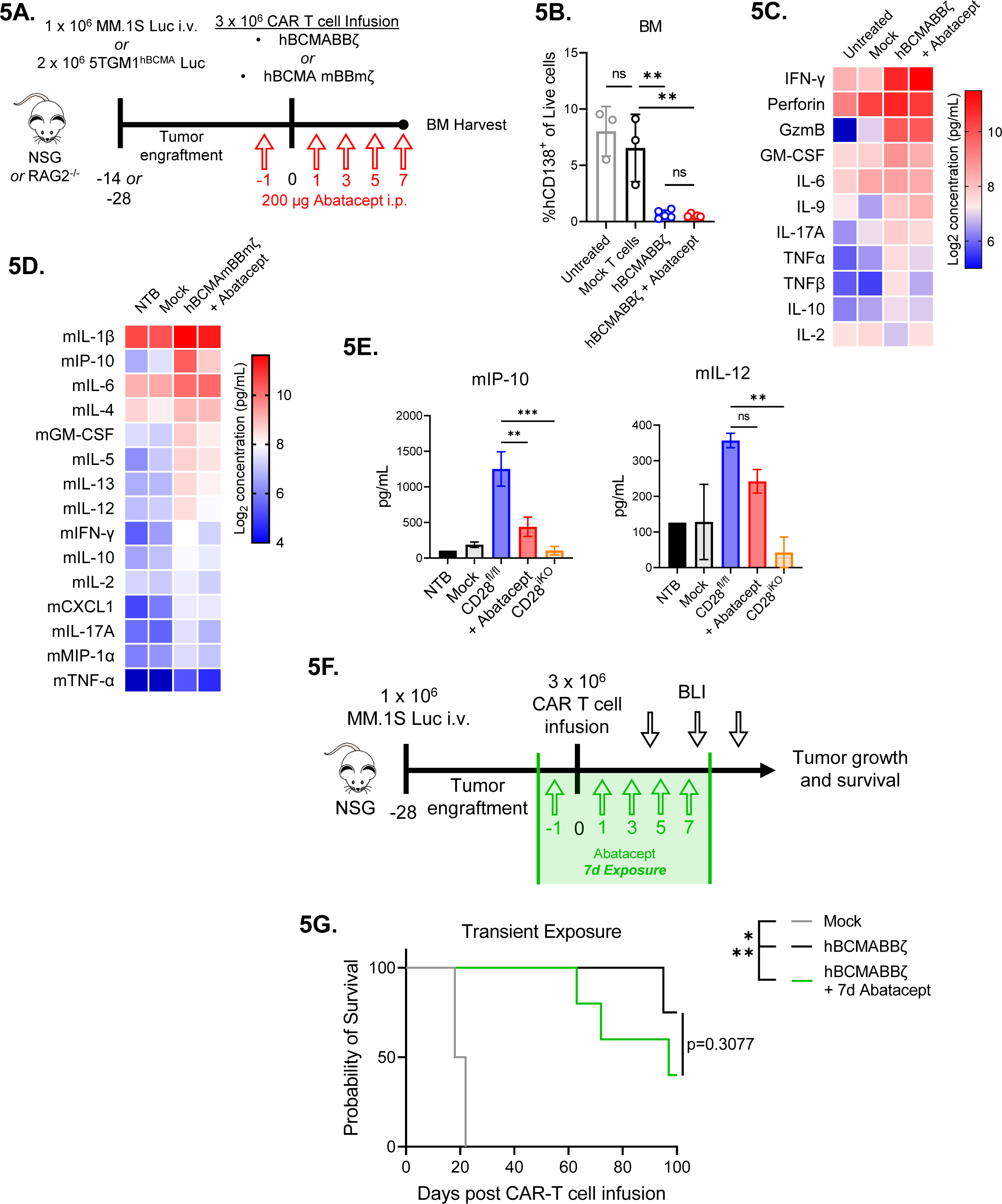
Transient CD28 blockade limits inflammatory cytokines in the MM BME without affecting survival of hBCMAmBBmζ CAR T cell treated mice. **(A)** Diagram of experimental setup. MM.1S bearing NSG mice were treated with hBCMABBζ CAR T cells ± 200μg abatacept on days −1, 1, 3, 5, 7 (5B and 5C) or 5TGM1^hBCMA^ bearing RAG2^-/-^ mice were treated with CD28^fl/fl^ or CD28^iKO^ hBCMAmBBmζ CAR T cells (5D and 5E) and euthanized 7 days post adoptive transfer for hind limb BM harvest. **(B)** Myeloma burden assessed by flow cytometry for human CD138^+^ cells in the bone marrow of MM.1S bearing mice treated as described in 5A. Bars represent mean ± SD, dots indicate individual mice. **p<0.01 by one-way ANOVA, ns = not significant. **(C)** Heatmap representation of human cytokine levels in the MM BME 7 days after treatment of MM.1S bearing mice with hBCMABBζ CAR T cells ± abatacept. Bilateral hind limbs were harvested, and BM was flushed into 15 µL PBS for multiplexed cytokine analysis. Log_2_ transformed cytokine concentrations represent the mean of 3-6 mice per group. **(D)** Heatmap representation of murine cytokine levels in the MM BME 7 days after treatment of 5TGM1^hBCMA^ bearing mice with hBCMAmBBmζ CAR T cells ± abatacept. Bilateral hind limbs were harvested, and BM was flushed into 15 µL PBS for multiplexed cytokine analysis. Log_2_ transformed cytokine concentrations represent the mean of 2-3 mice per group. **(E)** Bar graphs showing concentrations of murine IP-10 (left) and IL-12 (right) in the MM BME 7 days after treatment of 5TGM1^hBCMA^ bearing mice with CD28^fl/fl^ hBCMAmBBmζ CAR T cells ± abatacept versus CD28^iKO^ hBCMAmBBmζ CAR T cells. Bars represent mean ± SD of 2-3 biological replicates. **p<0.01, ***p<0.001 by one-way ANOVA. **(F)** Diagram of experimental setup. MM.1S bearing NSG mice were treated with hBCMABBζ CAR T cells ± 200μg abatacept on days −1, 1, 3, 5, 7 and tumor burden was monitored by bioluminescent imaging 2x/week until endpoint. **(G)** Kaplan – Meier analysis of survival of hBCMABBζ CAR T ± transient abatacept or mock transduced T cell treated MM.1S-luc bearing mice (n = 4-5 mice per CAR T cell treated group). Median survival of hBCMABBζ CAR T treated mice was >100 days post CAR T cell infusion vs. 97 days for hBCMABBζ CAR T + transient abatacept treated mice. Statistical significance was determined by log-rank Mantel-Cox test.

Somewhat surprisingly, inflammatory cytokine levels in the MM BME were dramatically reduced by CD28 deletion from mouse hBCMAmBBmζ CAR T cells (Fig. 2I) yet unaffected by CD28 blockade using abatacept in human hBCMABBζ CAR T cell treated mice (Fig. 5C). Such divergent findings may be due to contributions of cells other than CAR T cells to the mouse MM BME cytokine milieu or differences between how blockade of CD28 with abatacept and deletion of CD28 affect 4-1BB co-stimulated CAR T cells. To address these possibilities, we measured murine inflammatory cytokine levels in the MM BME of mouse hBCMAmBBmζ CAR T cell treated 5TGM1^hBCMA^ bearing mice. Abatacept diminished levels of murine inflammatory cytokines in the MM BME of hBCMAmBBmζ CAR T cell treated mice (Fig. 5D), yet not to the extent of CD28 deletion.

Notable among inflammatory cytokines affected by both abatacept treatment and CD28 deletion were IP-10, which is secreted by monocytes and stromal cells in response to IFN-γ^74^, and IL-12, which is mainly secreted by monocytes, macrophages, neutrophils, and dendritic cells^75^. It is therefore likely that other cells contribute to murine inflammatory cytokine production in the MM BME of hBCMAmBBmζ CAR T cell treated mice and given the magnitude of cytokine changes, that there is also a difference between how abatacept treatment and deletion of CD28 affects cytokine production.

### Transient CD28 blockade does not inhibit hBCMABB***ζ*** anti-MM activity

Based on our observation that abatacept could limit pro-inflammatory cytokine release but not early anti-tumor activity of hBCMABBζ CAR T cells, we predicted that transient abatacept exposure would not impair survival of myeloma bearing mice. To test this, abatacept was administered to MM.1S tumor bearing NSG mice from day −1 to day 7 post hBCMABBζ CAR T cell infusion (Fig. 5F). Transient abatacept exposure resulted in a slight reduction in tumor regression induced by hBCMABBζ CAR T cells (Supplemental Fig. 4C) but did not affect the long-term survival of hBCMABBζ CAR T cell treated myeloma bearing mice (Fig. 5H).

## Discussion

CAR T cell therapies targeting BCMA have shown curative potential in patients with relapsed/refractory multiple myeloma (MM)^25,27^. However, achieving long-term remissions remains an ongoing challenge, with one-quarter to greater than one-half of patients experiencing myeloma relapse within one year of CAR T infusion. In this study we set out to determine whether CD28 blockade using abatacept could sensitize MM cells to CAR T cell therapy in a manner analogous to standard chemotherapy^45,46,50^. In contrast to expectations, we found that abatacept limited efficacy of clinically relevant BCMA targeted, 4-1BB co-stimulated CAR T cells in an established human xenograft myeloma mouse model. Using a novel CD28 inducible knockout mouse model to generate CD28-deficient (CD28^iKO^) CAR T cells, we further revealed a previously unrecognized role for the endogenous CD28 receptor on 4-1BB co-stimulated CAR T cells. CD28 deletion did not alter BCMA targeted, 4-1BB co-stimulated CAR T cell cytotoxic capabilities nor alter inflammatory cytokine production *in vitro*, but rather resulted in diminished mitochondrial metabolism and a lower ratio of the oxidized form of nicotinamide adenine dinucleotide (NADH) to its reduced form (NAD^+^). These metabolic changes were associated with limited *in vivo* expansion of CD28^iKO^ 4-1BB co-stimulated CAR T cells and a reduction in CAR T cell induced inflammatory cytokine production in the multiple myeloma bone marrow microenvironment (MM BME). Abatacept treatment similarly reduced *in vivo* 4-1BB co-stimulated CAR T cell expansion and inflammatory cytokine production. Importantly however, short-term blockade of endogenous CD28 using abatacept during the first week following CAR T cell infusion reduced inflammatory cytokine levels in the MM BME without altering long-term survival of BCMA targeted, 4-1BB co-stimulated CAR T cell treated myeloma bearing mice.

Robust CAR T cell activation and expansion can induce systemic toxicities, including cytokine release syndrome (CRS), immune effector cell-associated neurotoxicity syndrome (ICANS), and immune effector cell-associated hematologic toxicity (ICAHT)^76–78^. In pivotal CAR T trials in MM^25,27^, 76 – 84% of patients experienced CRS, 18 – 21% experienced ICANS, and nearly all patients experienced hematologic toxicity, although these were generally transient. Current treatment options for CRS and ICANS include IL-6 receptor blockade with tocilizumab, corticosteroids for tocilizumab refractory cases, and the anti-IL-6 antibody siltuximab for tocilizumab and corticosteroid refractory toxicities^79^. Additionally, the IL-1 receptor antagonist anakinra is being explored as a potential prophylactic treatment to prevent CRS and ICANS^80^. Data presented here raise the possibility that abatacept (CTLA4-Ig), which is FDA approved for the treatment of rheumatoid arthritis, juvenile idiopathic arthritis, and psoriatic arthritis along with prevention of acute graft versus host disease^81,82^, may also be useful as prophylactic treatment to prevent toxicities brought on by 4-1BB co-stimulated CAR T cells. Whether abatacept could have similar utility in preventing CD28 co-stimulated CAR T cell toxicities is an open question currently lacking clinical relevance in the setting of multiple myeloma, where both FDA approved CAR designs contain a 4-1BB co-stimulatory domain.

Co-stimulation has long been known to be critical for anti-tumor effects of CAR T cells^83,84^, with different CAR-encoded co-stimulatory domains having distinct effects on CAR T cell properties^17^. Clinically available CARs contain either a CD28 or a 4-1BB co-stimulatory domain. CD28 co-stimulated CAR T cells exhibit rapid anti-tumor effector function but lack functional persistence associated with 4-1BB co-stimulated CAR T cells. Modulation of CAR-encoded CD28 signaling has resulted in improved functional persistence and reduced CAR T cell exhaustion in pre-clinical models^85,86^. Recent studies have hinted at a role for endogenous CD28 in determining CAR T cell efficacy. However, evidence for endogenous CD28 modulation of CAR T cell function was either indirect, in the case of CTLA4 knockout^55^, or complicated by co-expression of IL-12 from a fourth-generation armored CAR construct^87^. Data presented here provide the first direct evidence that endogenous CD28 affects efficacy of second-generation, 4-1BB co-stimulated CAR T cells comparable to those used to treat myeloma patients. These data raise important questions about how signaling from CAR co-stimulatory domains interfaces with signaling from endogenous co-stimulatory, and/or co-inhibitory receptors.

In light of recent evidence that CD28 co-stimulation in the tumor microenvironment is critical for effector differentiation and anti-tumor function of cytotoxic T cells^56,88^, the context in which endogenous CD28 expressed on 4-1BB co-stimulated CAR T cells encounters ligands may influence anti-tumor efficacy and/or inflammatory cytokine production. B lineage tumors targeted clinically by CAR T cells are characterized by high levels of CD28 ligand expression, with CD80, CD86, or both expressed on tumor cells in more than half of myeloma, non-Hodgkin’s lymphoma, and B-ALL paitents^89–91^. CD80 and CD86 are also expressed on antigen presenting cells throughout the body of cancer patients and may have similar or disparate effects on CAR T cells, likely based on whether CAR engagement occurs concurrently with CD28 engagement. If future studies find that the context in which endogenous CD28 is engaged matters, CD80 and/or CD86 expression patterns may become useful in determining whether CD28 or 4-1BB co-stimulated CAR T cells are used to treat particular patients.

Overall, results presented here provide the first direct evidence that endogenous CD28 is important for driving anti-tumor function of 4-1BB co-stimulated CAR T cells. These results raise many interesting and important biological questions about co-stimulation and signaling in CAR T cells and, perhaps more importantly raise the possibility that blocking endogenous CD28 signaling may abrogate some of the toxic side effects associated with CAR T therapy. Future studies aimed at optimizing methods and timing of CD28 blockade have the potential to lead to improved clinical strategies for limit toxicities while maintaining CAR T cell efficacy.

## Acknowledgements

The authors would like to thank Steven Turowski and Joseph Spernyak of the Translational Imaging Shared Resource (TISR) at Roswell Park for their assistance with bioluminescent imaging studies as well as the Cancer Center Support Grant: P30CA016056 and shared instrumentation grant: S10OD16450. We would also like to acknowledge Courtney Ryan for her technical expertise and operation of the Seahorse XFe96 extracellular flux analyzer within the Immune Analysis Facility at Roswell Park. Finally, we would like to thank Bristol Myers Squibb (BMS) for providing abatacept for use in our studies under the non-sponsored research agreement IM101-902.

## Author Contributions

**Conception and design:** M. Lieberman, K. Lee, S. Olejniczak

**Development of methodology:** M. Lieberman, K. Lee, S. Olejniczak

**Acquisition of data (provided animals, managed studies, provided facilities, etc.):** M. Lieberman, J. Bramson, R. Brentjens, K. Lee, S. Olejniczak

**Analysis and interpretation of data:** M. Lieberman, K. Lee, S. Olejniczak

**Writing, review, and/or revision of the manuscript:** M. Lieberman, K. Lee, S. Olejniczak

**Administrative, technical, or material support:** M. Lieberman, J. Tong, N. Odukwe, T. Purdon, R. Burchett, A.J.R. McGray, B. Gillard, C. Brackett, C. Chavel

**Study supervision:** S. Olejniczak

All authors have read and agreed to the published version of the manuscript.

## Institutional Review Board Statement

This study involves de-identified healthy donor samples collected under an approved protocol reviewed by the Roswell Park Institutional Review Board (IRB): BDR 115919. All animal experiments were performed in accordance with the Institutional Animal Care and Use Committee (IACUC) guidelines and were approved under experimental IACUC protocol: 1094M.

## Data Availability Statement

All data associated with this paper are included in the manuscript and supplementary materials. Requests for resources and reagents should be directed to and will be fulfilled by the corresponding author, Scott H. Olejniczak

## Supplemental Figure Legends

**Supplemental Fig. 1: Characterization of human hBCMABB**ζ **CAR T cells and myeloma target cells.**

**(A)** CD4:CD8 ratio in CAR T cell infusion products generated from PBMCs of healthy donors. Relative frequency of CD4^+^ and CD8^+^ CAR T cells assessed by flow cytometry. Each data point represents an independent donor (n = 8).

**(B)** Representative flow cytometry dot plot depicting the frequency of CAR transduced cells.

**(C)** Transduction efficiency (left) and mean fluorescence intensity MFI; right) of hBCMABBζ CAR expression on human CD3^+^ T cells determined using αG4S linker antibody and flow cytometry. Dots represent independent donors (n = 10).

**(D)** Surface CD28 protein expression on CD4^+^ and CD8^+^ hBCMABBζ CAR T cells determined by MFI. Each data point represents an independent donor (n = 6). **p<0.01 by unpaired Student’s t test **p<0.01.

**(E)** Flow cytometry histograms depicting surface expression of MM defining phenotypic markers CD138 and BCMA along with co-receptors CD28, CD80, and CD86 on human MM cell lines, MM.1S and U266.

**(F)** Heatmap representation of culture supernatant human cytokine concentrations measured by multiplexed Luminex assays at the conclusion of a 24-hr. co-culture of hBCMABBζ CAR T cells ± abatacept with MM.1S myeloma cells. Log_2_ transformed cytokine concentrations represent the mean of 5 independent experiments using CAR T cells generated from 5 healthy donors.

**Supplemental Fig. 2: Characterization of mouse CD28^iKO^ hBCMAmBBmζ CAR T cells and myeloma target cells.**

**(A)** CD4:CD8 ratio in CAR T cell infusion products generated from CD28^fl/fl^ or CD28^iKO^ splenocytes. Relative frequency of CD4^+^ and CD8^+^ CAR T cells assessed by flow cytometry. Each data point represents an independent experiment (n = 6 per group).

**(B)** Representative flow cytometry histograms depicting CD28^fl/fl^ and CD28^iKO^ hBCMAmBBmζ CAR transduced T cells.

**(C)** Transduction efficiency of the hBCMAmBBmζ CAR into CD28^fl/fl^ versus CD28^iKO^ mouse T cells, expressed as percentage of CD3^+^ T cells determined using αG4S linker antibody and flow cytometry. Dots represent independent experiments (n ≥ 10 per group).

**(D)** Schematic of lentiviral vector used to transduce 5TGM1 cells to express human BCMA (hBCMA) and firefly luciferase.

**(E)** Flow cytometry histogram depicting surface hBCMA expression on transduced and sorted 5TGM1 mouse myeloma cells.

**Supplemental Fig. 3:**

**(A)** CD28^fl/fl^ and CD28^iKO^ mBBmζ CAR T cells assessed by MFI of MitoGreen staining.

**Supplemental Fig. 3: Short duration of CD28 blockade does not impair early anti-tumor responses but may reduce systemic toxicities by dampening pro-inflammatory cytokines in the BM, related to figure 4 and figure 5**.

**(A)** Flow cytometry gating strategy used to determine tumor burden and CAR T cell frequency following BM harvest of 5TGM1 bearing RAG2^-/-^ mice treated with CD28^fl/fl^ mBBmζ CAR T cells ± abatacept for 7 days.

**(A)** Tumor burden assessed by the frequency of mCD138^+^ B220^-^ cells in the BM of 5TGM1 bearing RAG2^-/-^ mice treated with CD28^fl/fl^ mBBmζ CAR T cells ± abatacept for 7 days. Each data point represents an individual tumor bearing host. Significance determined by one-way ANOVA.

## Supplemental Table Captions

1. Mouse and human antibodies used for cell phenotyping analyses performed by flow cytometry.

2. Mouse primer sequences used for qRT-PCR.

