## Supplementary figures and images for "Endogenous CD28 drives CAR T cell responses in multiple myeloma"

### Supplemental Figures

##### *Supplemental Fig. 1:*

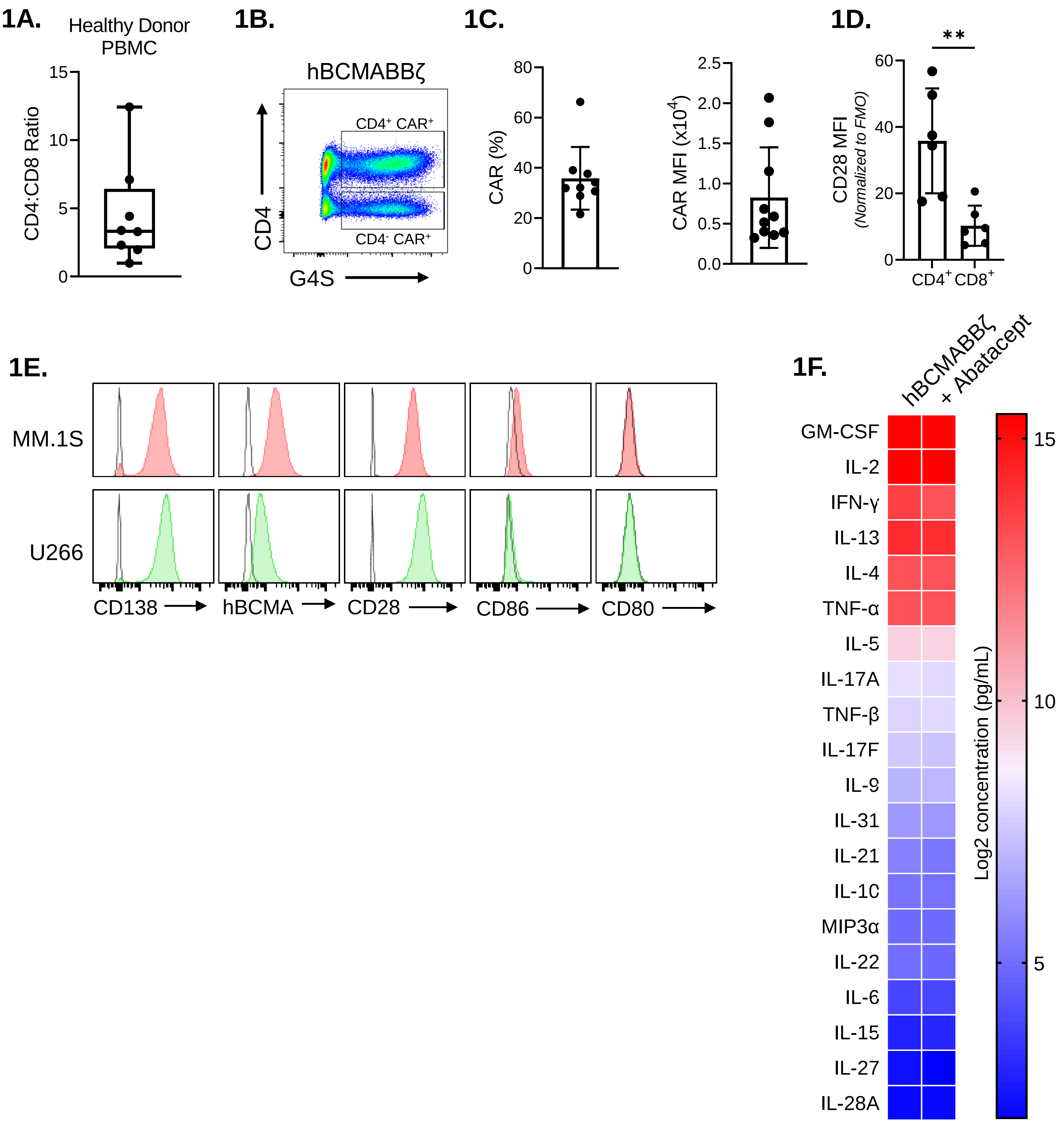

#### Supplemental Fig. 2:

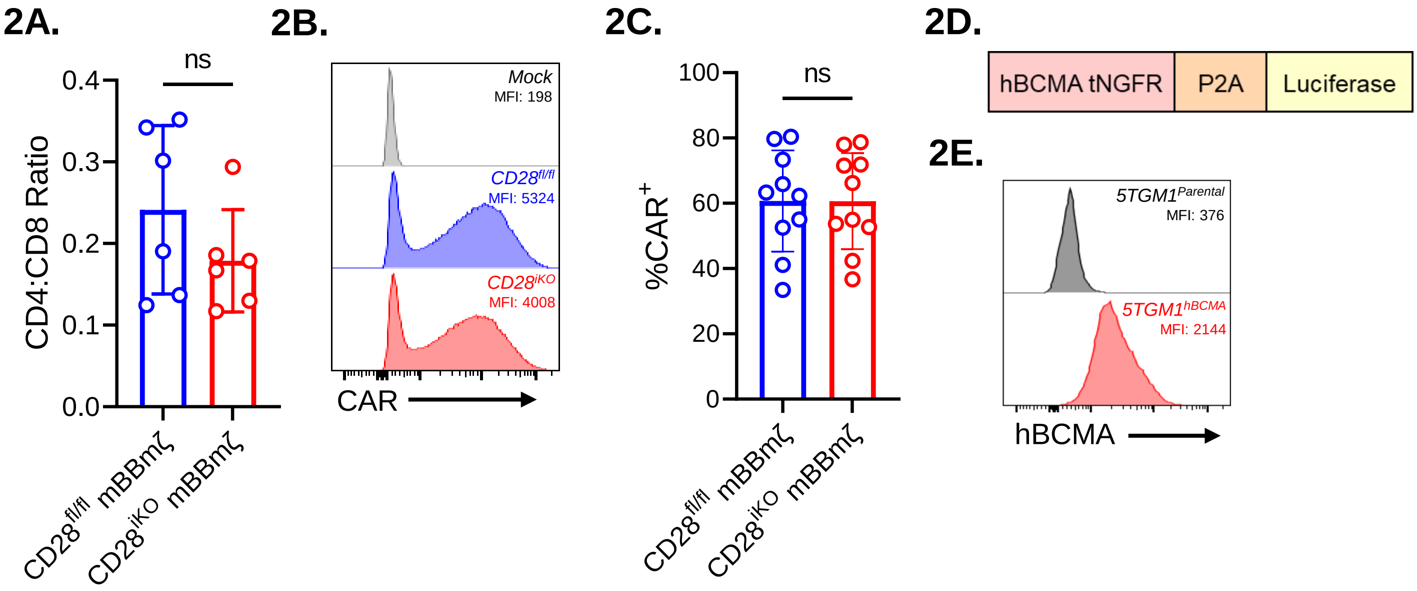

***Supplemental Fig. 3:***

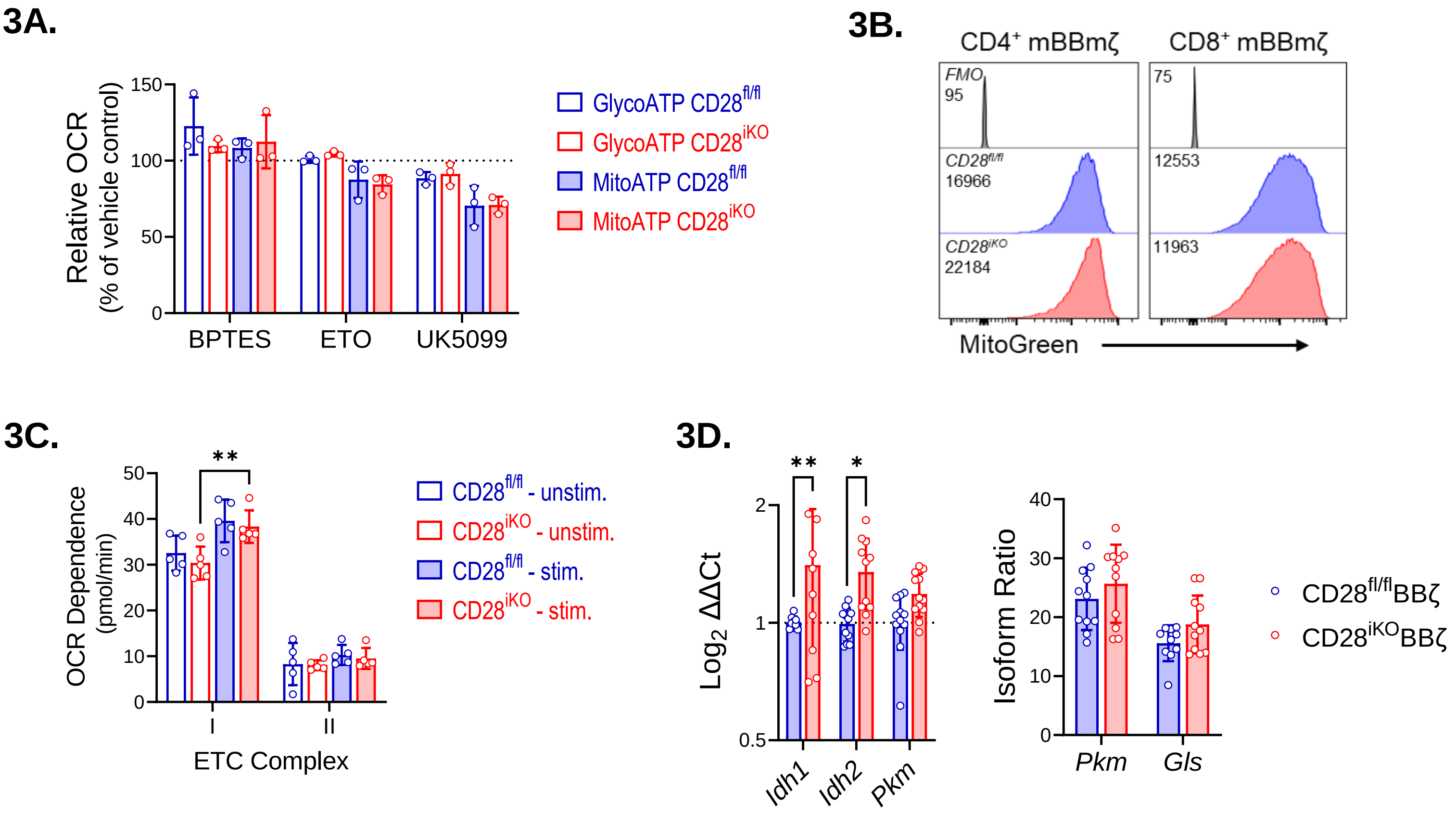

##### *Supplemental Fig. 4:*

### 4A.

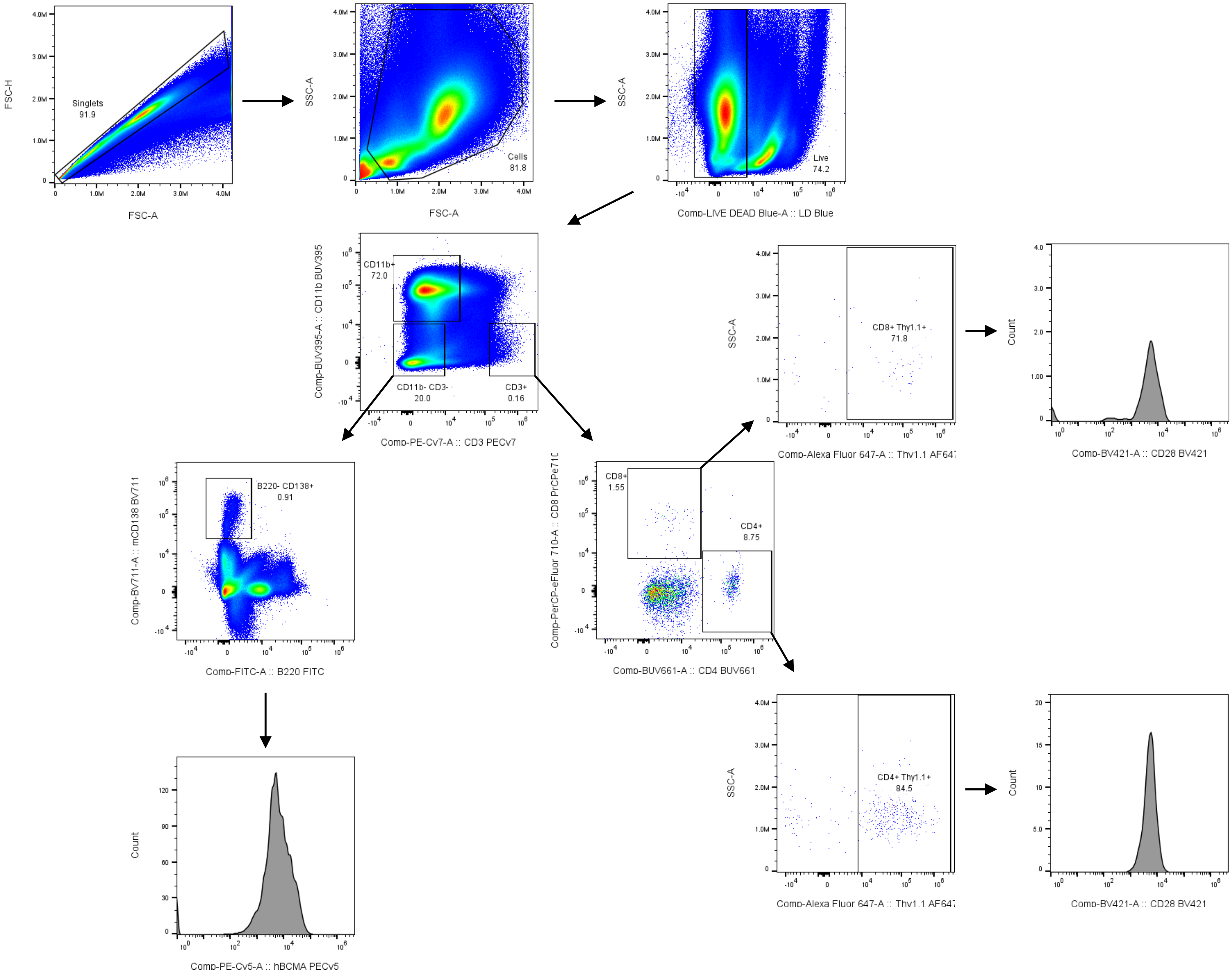


### 4B.

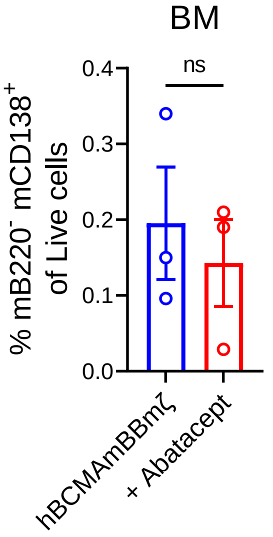


### 4C.

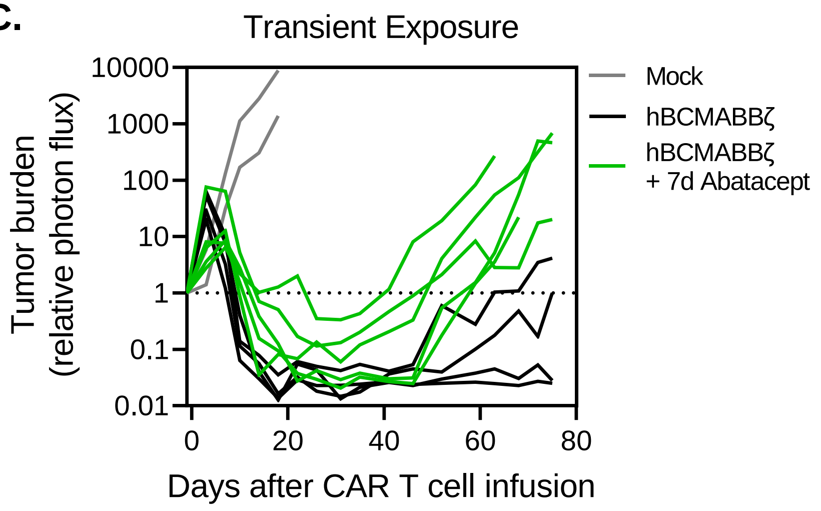
